# A robust nonlinear low-dimensional manifold for single cell RNA-seq data

**DOI:** 10.1101/443044

**Authors:** Archit Verma, Barbara E. Engelhardt

**Affiliations:** Department of Chemical and Biological Engineering, Princeton University, Princeton, NJ 08544, USA; Department of Computer Science, Center for Statistics and Machine Learning, Princeton University, Princeton, NJ 08540, USA

**Keywords:** Manifold Learning, Gaussian Process Latent Variable Model, cell types, Single Cell RNA-Sequencing (scRNA-seq), Robust Statistics

## Abstract

Modern developments in single cell sequencing technologies enable broad insights into cellular state. Single cell RNA sequencing (scRNA-seq) can be used to explore cell types, states, and developmental trajectories to broaden understanding of cell heterogeneity in tissues and organs. Analysis of these sparse, high-dimensional experimental results requires dimension reduction. Several methods have been developed to estimate low-dimensional embeddings for filtered and normalized single cell data. However, methods have yet to be developed for unfiltered and unnormalized count data. We present a nonlinear latent variable model with robust, heavy-tailed error and adaptive kernel learning to estimate low-dimensional nonlinear structure in scRNA-seq data. Gene expression in a single cell is modeled as a noisy draw from a Gaussian process in high dimensions from low-dimensional latent positions. This model is called the Gaussian process latent variable model (GPLVM). We model residual errors with a heavy-tailed Student’s t-distribution to estimate a manifold that is robust to technical and biological noise. We compare our approach to common dimension reduction tools to highlight our model’s ability to enable important downstream tasks, including clustering and inferring cell developmental trajectories, on available experimental data. We show that our robust nonlinear manifold is well suited for raw, unfiltered gene counts from high throughput sequencing technologies for visualization and exploration of cell states.

## 1. Background

High-throughput single cell RNA sequencing (scRNA-seq) is a powerful tool for cataloguing cell types and cell states, and investigating changes in expression over cell developmental trajectories. Droplet-based methods encapsulate individual cells with unique barcode tags that are ligated to cellular RNA fragments (Zheng et al., 2017). Sequenced reads are mapped to both a gene and a cell, creating a high-dimensional cell by gene count matrix–with hundreds to millions of cells and twenty thousand genes per human cell. These cell by gene count matrices contain a substantial proportion of zeros because of low coverage sequencing per cell (i.e., *dropout*), and also contain substantial variance from both technical and biological sources of noise (Buettner et al.). Computational tools for analyzing scRNA-seq results thus require initial dimension reduction to a lower dimensional manifold capturing gene expression patterns for regularization and computational efficiency. Dimension reduction techniques are conducive to noise reduction (Buettner et al.; Eraslan et al., 2018), sub-population identification (Pierson and Yau; Haghverdi et al., 2016), visualization (Amodio et al.; Van Der Maaten and Hinton), pseudotemporal ordering of development stages (Ahmed et al., 2018; Trapnell et al.; Lönnberg et al., 2017), and imputation (Li and Li, 2018). Lower-dimensional mappings also provide convenient visualizations that inform analytic methods and future experiments.

Linear dimension reduction techniques are commonly used as a first step to downstream analyses. Principal component analysis (Hotelling, 1933) (PCA) – the projection of a high dimensional space onto orthogonal bases that capture the directions of greatest variance – is the first step of several scRNA-seq analysis packages such as PAGODA (Fan et al., 2016) and Waterfall (Shin et al.). Zero-Inflated Factor Analysis (Pierson and Yau) (ZIFA) extends the factor analysis (Harman, 1960) paradigm of a linear mapping onto low dimensional latent dimensions to allow dropouts modeled by Bernoulli random variables to account for the excess of zero counts in scRNA-seq data. Independent component analysis (Comon, 1994) (ICA), which assumes non-Gaussian observations, and canonical correlation analysis (Hotelling, 1936) (CCA), which allows for multiple observations, have also been used as dimension reduction techniques for studying cell developmental trajectories (Trapnell et al.) and for experimental batch correction (Butler et al.).

More sophisticated models eschew the linearity assumption to find richer nonlinear structure in the data. The t-distributed Stochastic Neighbors Embedding (t-SNE) (Van Der Maaten and Hinton) is a popular visualization tool. t-SNE computes the similarity between two points in high dimensional space with respect to a Gaussian kernel distance metric, and estimates a lower dimensional mapping with similarity with respect to a Student’s t-distribution metric that minimizes the Kullback-Leibler divergence between the similarity distributions in high and low dimensions. The Gaussian kernel in t-SNE includes a perplexity parameter that controls the decay rate of similarity across the distance between cells. Diffusion maps, used in packages such as Destiny (Angerer et al., 2016), are another tool for nonlinear low dimensional mapping that perform linear decomposition on a kernel similarity matrix of high dimensional observations. SAUCIE (Amodio et al.) implements a variational autoencoder, or a deep neural network that compresses data with the goal of creating an optimal reconstruction from the compressed representation, to execute several single cell tasks. Similarly, scVI (Lopez et al.) uses deep neural networks to create a probabilistic representation in latent space for batch correction, visualization, clustering, and differential expression. One Bayesian probabilistic technique is the Gaussian process latent variable model (Titsias and Lawrence; Lawrence) (GPLVM), which used by scLVM Buetner et al.) for noise reduction, and GPfates (Lönnberg et al., 2017) and GrandPrix (Ahmed et al., 2018) for pseudotemporal ordering. The GPLVM models observations (i.e., cells) as draws from Gaussian processes representations of lower dimensional latent variables.

While current methods for dimension reduction have been successful with early sequencing experiments and filtered expression data, they are limited in their capacity to accurately represent and inform analyses of raw, high throughput sequencing experiments. Linear methods such as PCA and ZIFA are ill-suited for capturing highly nonlinear biological processes across developmental phase, and many implementations scale poorly with increased sample size. Current non-linear methods are highly sensitive parameter choices, including perplexity for t-SNE, kernel variables for diffusion maps, and network architecture for VAEs. Latent dimensions of t-SNE have no global structure, making embedded positions difficult to interpret and leading to uninformative mappings beyond two dimensions. Downstream analyses of t-SNE results are hindered by an inability to map back to observation space. VAEs, like most neural networks, require tens to hundreds of thousands of cells for accurate estimation, which may not be available in smaller experiments. Current methods, particularly those using the GPLVM, work only with filtered, normalized data and incorporate prior information to facilitate the latent mapping.

Robust statistical methods are a natural solution to modeling noisy, sparse count data. We introduce the t-Distributed Gaussian Process Latent Variable Model (tGPLVM) for learning a low dimensional embedding of unfiltered count data. We introduce three features to the basic GPLVM: 1) a robust Student’s t-Distribution noise model; 2) a weighted sum of non-smooth covariance kernel functions with parameters estimate from the data; 3) sparse kernel structure. The heavy tailed Student’s t-Distribution improves robustness to outliers, previously demonstrated in Gaussian process regression (Tang et al.; Vanhatalo et al.). Matérn kernels have been successfully used in time series modeling to capture non-smooth trajectories (Ahmed et al., 2018). The sparse kernel structure allows us to effectively reduce the number latent dimensions based on the actual complexity of the data. Our implementation of tGPLVM accepts sparse inputs produced from high-throughput experimental cell by gene count matrices.

We demonstrate tGPLVM’s ability to estimate informative manifolds from noisy, raw single cell count matrices and highlight its applicability to multiple downstream tasks. We show improved cell type identification via clustering on the estimated latent space using a data set of cerebral cortex cells labeled with estimated cell type (Pollen et al.). We find that the tGPLVM manifold can learn pseudotemporal ordering from a batch of *Plasmodium*-infected mouse cells sequenced across time post exposure (Lönnberg etal., 2017). Finally, we demonstrate that tGPLVM can be used on unprocessed, unnormalized count data from recent high-throughput sequencing methods (Zheng et al., 2017) and used to explore gene expression across cell states. We implement a scalable inference algorithm that can fit hundreds of thousands to millions of cells.

## Results

The t-distributed Gaussian process latent variable model is a nonlinear latent variable model that captures high dimensional observations in low dimensional nonlinear latent space. Expression of each of *P* genes across all cells is modeled as a draw from a multivariate normal distribution with the covariance a function of the low-dimensional, latent positions. The observation error is modeled with a heavy tailed Student’s t-Distribution with four degrees of freedom to robustly account for variance in the count data due to technical and biological noise relative to a normally distributed error model (O’Hagan, 1979, 1988). Here, we use a weighted sum of Matérn/2, Matérn 3/2, Matérn 5/2, and squared exponential kernel functions to model non-smooth manifolds.

### Nonparametric manifold learning improves cell type identification

First, we evaluated the ability of tGPLVM and commonly used single cell dimension reduction methods to distinguish distinct cell types. tGPLVM, PCA, ZIFA, and t-SNE were used to map cells labeled with their inferred cell type from the Pollen data (Pollen et al.) to latent spaces varying from two to nine dimensions. With more than two latent dimensions, tGPLVM produced clusters that best corresponded to the actual cell type labels of the four methods (Figure 2). Including, Matérn, kernels in tGPLVM improves cell type separation in the latent space as measured by normalized mutual information and adjusted rand score (Supplementary Figure 1). Inclusion of Matérn kernels also reduces the uncertainty of posterior estimates of the latent embedding as measured by the average scale parameter of the latent position (Supplementary Figure 2). These results suggest that a robust Bayesian nonparametric manifold is superior to current dimension reduction algorithms for identifying and visualizing distinct cell types captured by scRNA-seq experiments.

**Figure 1:**
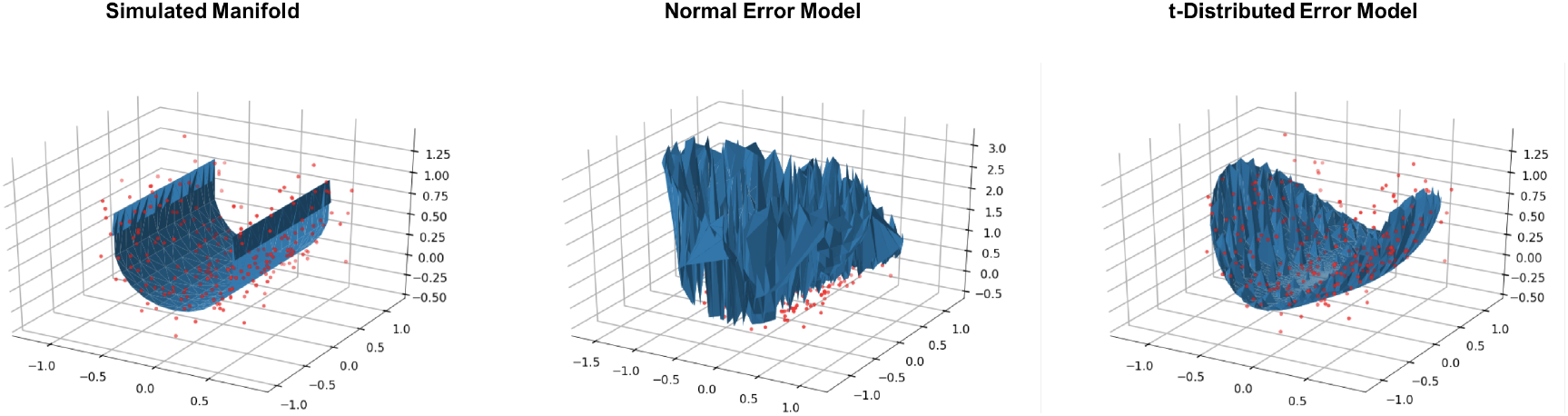
Comparison of error models on the same observations (red) in a multidimensional Gaussian process. Data are simulated from a smooth manifold with t-distributed error (left). A normally distributed error model (center) overfits to the data and fails to find the manifold structure due to the outliers as compared to the manifold estimated by a GP with t-distributed error (right).

**Figure 2:**
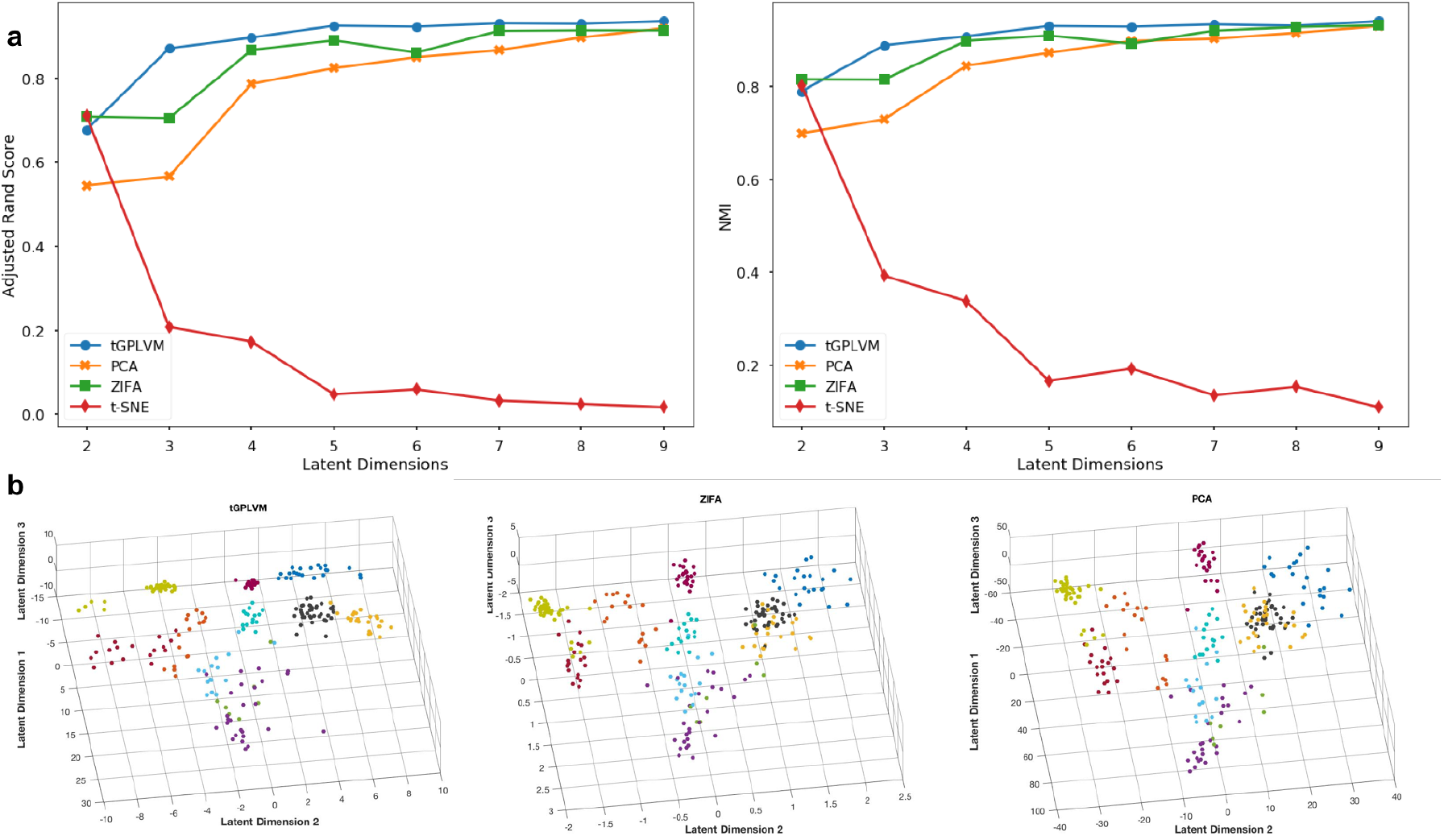
Comparison of manifold learning methods on 11 neural and blood cell populations (Pollen et al.). (a) Average ARS (left) and NMI (right) of ten K-means cluster labels versus available cell type labels with respect to the number of latent dimensions. (b) Three dimensional latent mappings from tGPLVM, PCA, and ZIFA colored by inferred cell type label. t-SNE (not pictured) collapses in three dimensions.

### Nonparametric manifold learning can reconstruct development time scales without prior information

Next we test the flexibility of tGPLVM to continuous cellular developmental trajectories by fitting latent mappings for a batch of mouse Th1 and Tfh cells sequenced over seven days after infection with *Plasmodium* (Lönnberg etal., 2017). Visually, we find that the latent mapping from tGPLVM represents the temporal relationships accurately, with most cells positioned among cells from the same or adjacent time points. We build a minimum spanning tree on the latent mappings to infer developmental trajectories. For a two dimensional mapping, only tGPLVM accurately spans the first time point (day 0) to the final time point (day 7). PCA, ZIFA, and alternate GPLVM models with different error or kernel choices find endpoints of the tree in days 2 or 4. t-SNE is able to separate cells based on time but does not accurately reconstruct the ordering and is clearly sensitive to outliers (Figure 3). This suggests that tGPLVM is a superior dimension reduction technique for identifying developmental pathways in unlabeled settings.

**Figure 3:**
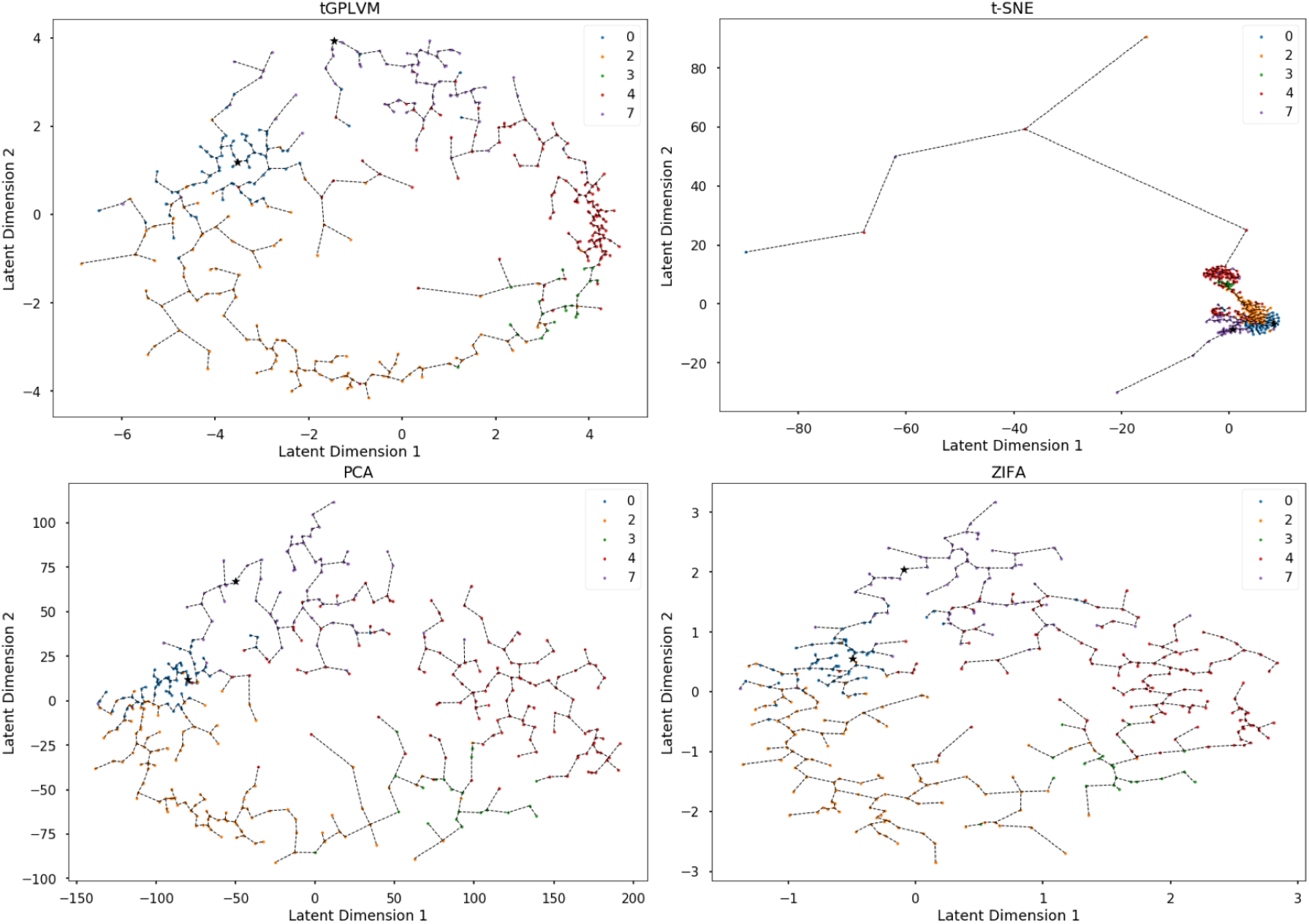
Comparison of manifold learning Methods on *Plasmodium*-infected Th1 and Tfh cells (Lönnberg etal., 2017): Plot of two dimensional latent mapping from tG-PLVM, PCA, ZIFA, and t-SNE. Labels indicate days after infection prior to sequencing. Dotted lines represent connections along the minimum spanning tree.

### Nonparametric manifold learning improves visualization of raw count data and captures cell state

Next, we tested tGPLVM’s performance on unfiltered count data. Models were fit on ~10,000 CD34+ peripheral blood mononuclear cells (PBMCs) sequenced on a high throughput parallel 10x system (Zheng et al., 2017). Each model was able to find three distinct regions based on expression patterns (Figure 4). PCA is dominated by total counts, with cells with more reads moving further away in latent space, and more frequent cell types dominate the space (Engelhardt and Stephens, 2010). CD34 is a marker for hematopoeitic stem cells (Sidney et al., 2014), which differentiate into myeloid and lymphoid cells. From tGPLVM, we can observe this separation from different expression patterns in progenitor cells across dimensions. Dimension three correlates with myeloid cells, demonstrated visually by marker *TYROBP* (Tomasello and Vivier, 2005) (Pearson’s *r* = 0.647; Figure 4), in addition to correlations with macrophage-associated genes (Donato et al.; Xia et al., 2018) *S100A4* (Pearson’s *r* = 0.623) and *S100A6* (Pearson’s *r* = 0.665). Dimension two correlates to lymphoid cells, visualized by marker *LTB* (Browning et al., 1993) (Pearson’s *r* = 0.306; Figure 4), and further supported by correlation with lymphocyte specific protein-1 *LSP1* (Pearsons’s *r* = 0.481). Dimension one corresponds to general cellular functions, with strong correlation with mitochondrial activity genes *COX5A* (Pearson’s *r* = 0.587) and *STOML2* (Pearson’s *r* = 0.474), and shown with *CLTA*, an endocytosis-mediating gene (Stelzer et al., 2016) (Pearson’s *r* = 0.461; Figure 4). These distinct expression patterns reflect the broadly different immune cellular functions into which hematopoietic stem cells may develop. Gradients of expression levels projected on tGPLVM embeddings can be used to further interrogate changes in cell states from different experiments and across these manifolds.

**Figure 4:**
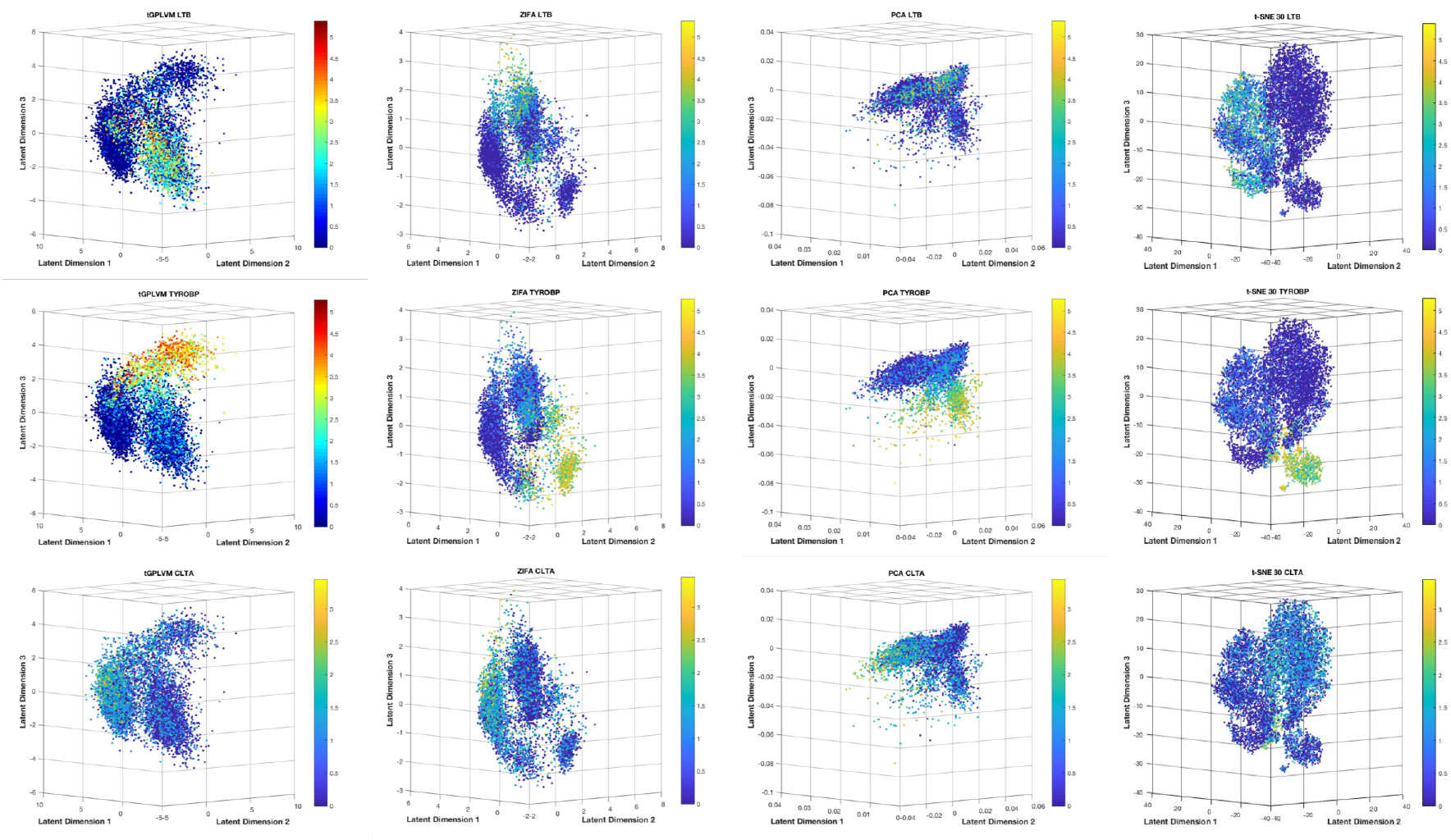
Manifold learning methods on unprocessed CD34+ PBMCs (Zheng et al., 2017) counts. tGPLVM shows the best separation of expression patterns based on cell state marker genes. Color bars indicate log2(1 + *Y*), where *Y* represents total counts per gene per cell.

### tGPLVM scales to a million cells

Finally, we evaluate the ability of tGPLVM and related methods to fit embeddings for unfiltered, unnormalized, high throughput scRNA-seq data. Models with two latent dimensions were fit on subsamples from 100 to 1 million cells from the 10x 1 million mouse brain cell data (Zheng et al., 2017). tGPLVM and PCA are the only methods that can fit one million cells in a computationally tractable way (Figure 5). ZIFA is slower than tGPLVM by an order of magnitude consistently across sample sizes. Since ZIFA requires a dense input, its input cell count matrix is limited in our framework to approximately 100,000 cells. While t-SNE’s implementation can input a sparse matrix format, it does not converge beyond 10^4^ samples.

**Figure 5:**
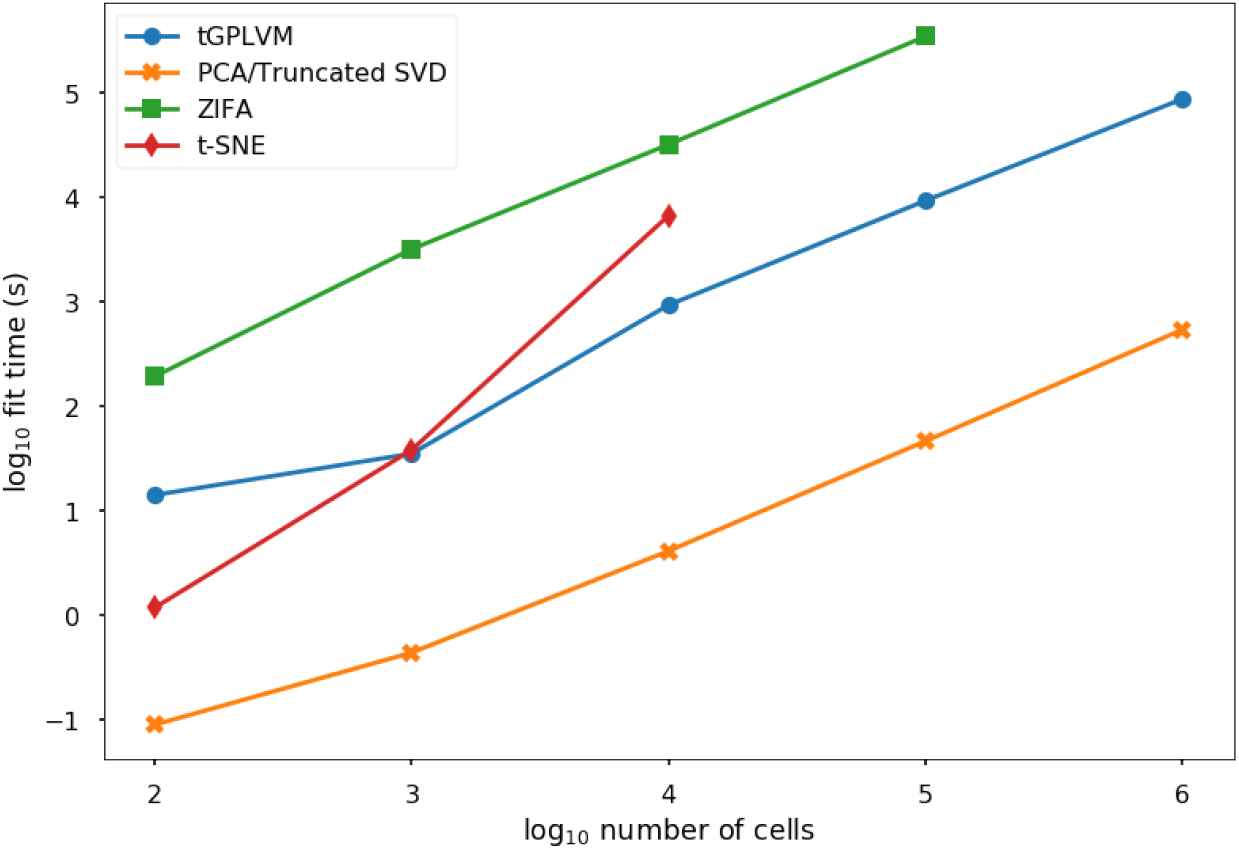
Time to fit a two dimensional embedding vs. sample size on a 16 VCPU, 224 GB memory high performance computing system.

To check that the embedding has biological significance, we again used Pearson’s correlation to identify genes whose expression is correlated with latent dimensions. We find that latent dimension one corresponds to increased expression of genes associated with the circulatory system and hemoglobin, such as *HBB-BS* (Pearson’s *r* = 0.320) and *HBA-A1* (Pearson’s *r* = 0.316; Figure 6b). Dimension two correlates with genes such as *TUBA1A* (Pearson’s *r* = 0.474) and *FEZ1* (Pearson’s *r* = 0.427) that are associated with neural cells (Figure 6a). The ability of tGPLVM to scale to high-throughput data and capture global structure from unnormalized count matrices makes it a powerful method for analyzing future single cell experiments.

**Figure 6:**
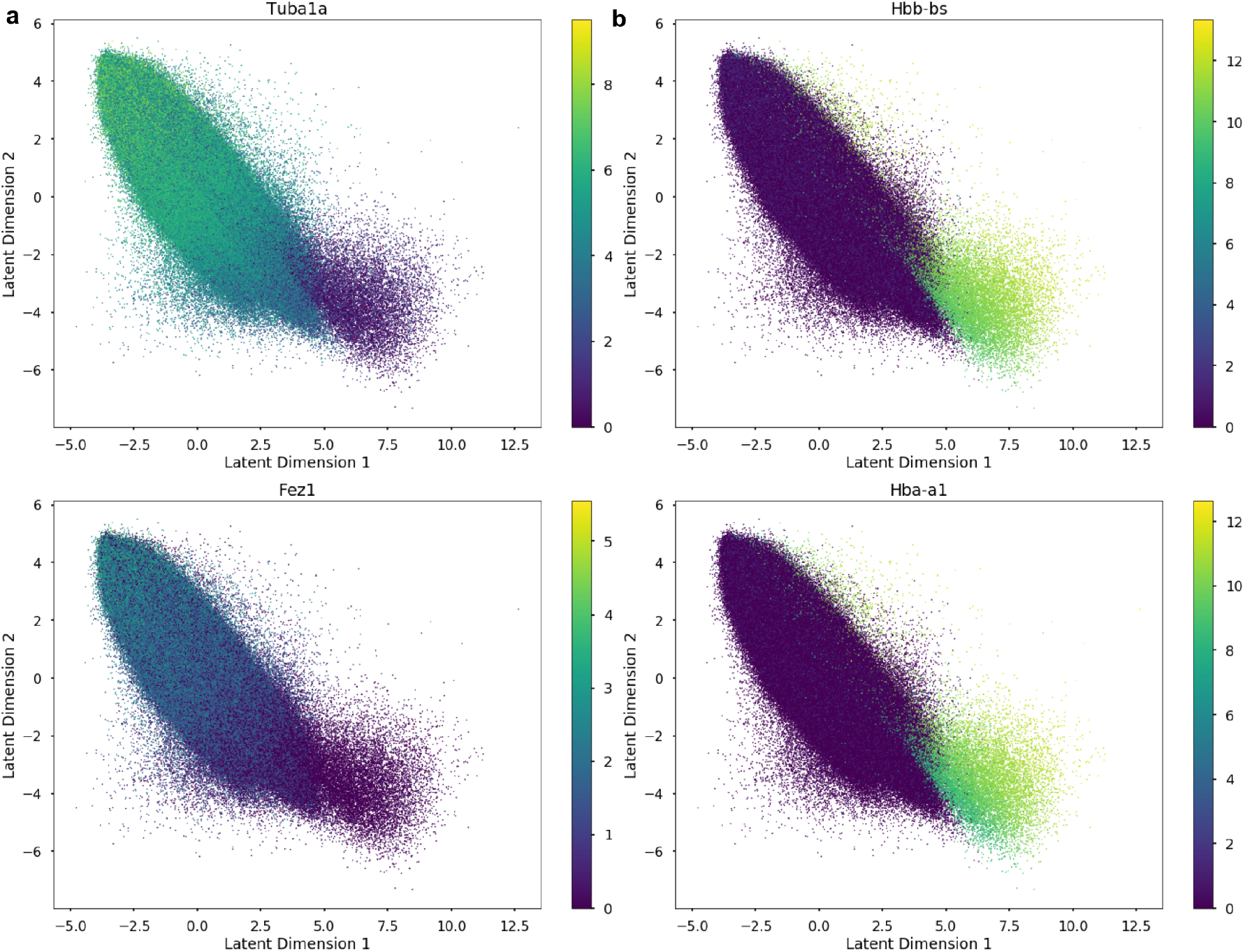
Gene expression patterns across tGPLVM manifold in 1 million mouse brain cells (Zheng et al., 2017). Latent dimension one separates circulatory and blood genes. Latent dimension two is correlated with neural genes. Color bars indicate log_2_(1 + *Y*), where *Y* represents total counts per gene per cell.

## Discussion

We present a Bayesian nonparametric model for robust nonlinear manifold estimation in scRNA-seq settings. tGPLVM captures transcriptional signals in single cell data using a robust Student’s t-distribution noise model and integrating adaptive kernel structure in settings with no *a priori* information about clusters or sequencing order. Our results show that tGPLVM is flexible to cell type, cell development, and cell perturbation experiments and can learn informative mappings from filtered and processed data as well as unfiltered raw count data. tGPLVM scales to the size of a million cells as produced by the latest single cell sequencing systems. Despite the sparsity, these data are complex and require several factors to capture variation; we did not use the ARD kernel parameters to remove dimensions for any of our experiments. However, the embedding dimensions in our experiments were able to capture informative representations of these complex data, and as the number of latent dimensions increased some would eventually be removed for being redundant in capturing these complex transcriptional profiles. We expect that the estimated latent mappings can be used for more sophisticated, nonparametric approaches for a variety of single cell tasks from normalization and imputation to cell type identification. We also hope that this robust manifold estimation can be used for other types of data with noisy outliers and sparse features.

## Methods

### The t-Distribution Gaussian process latent variable model

The tGPLVM assumes that samples in high dimensional space are noisy observations of a Gaussian process of lower dimensional latent features. Let *Y* ∈ ℝ^*N ×P*^ represent *N* observations in a high dimensional space of dimension *P*, and let *X* ∈ ℝ^*N ×Q*^ represent the same observations in a lower dimensional space *Q* ⪡ *P*. Each sample *x_n_* in *{n* ∈ 1, 2, …*, N }* is assumed to be drawn from a Q dimensional multivariate normal distribution with identity variance:

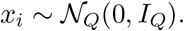

Noiseless observations of each of the *P* high dimensional features across *N* samples, *f_p_*(*X*), are draws from a zero-mean Gaussian process of *x* across a weighted sum of *M* kernels:

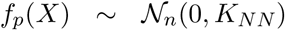

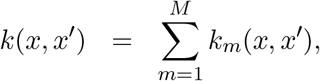

where *K_NN_* represents the *N × N* covariance matrix defined by *k*(*x, x*′). In the traditional GPLVM, observations *y_n,p_* are noisy realization of a Normal distribution with mean *f_n,p_* and variance *τ*^2^:

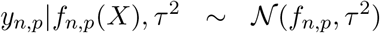

For tGPLVM, each observation *y_n,p_* is drawn from a heavy-tailed Student’s t-distribution with a set degrees of freedom *ν* and feature-specific variance 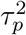 (Tang et al.):

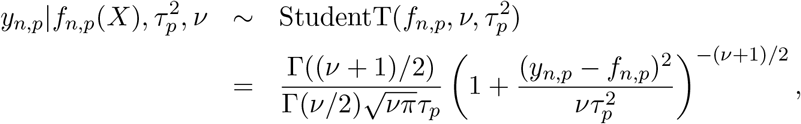

where we use *f_n,p_* to represent the *n*th component of the *N* dimensional vector *f_p_*(*X*). We set *ν* = 4 based on previous work with the supervised Gaussian process with t-distributed error (Tang et al.; Vanhatalo et al.).

The kernel we use is a flexible sum of an automatic relevance determination (ARD) squared exponential kernel and three different Matérn ARD kernels each with hyperparameters scale *σ_k_* and length scales 𝓁_*k,q*_. Each ARD dimension-specific length scale, *l_k,q_* indicates the distance of that latent dimension over which points are similar. Letting *r* represent the length scale-weighted distance in latent space, the kernels are defined as:

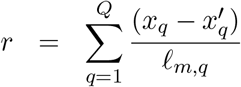

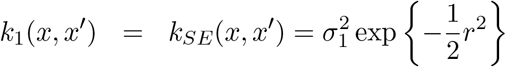

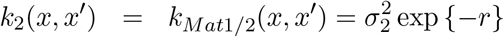

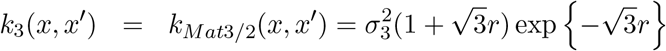

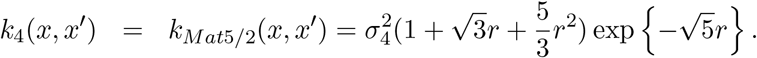

We use Black Box Variational Inference (Ranganath et al.) to estimate the posterior distribution for tGPLVM. We adapt the variational distributions from prior work (Damianou et al., 2016). Inference is implemented in Python using Edward (Tran et al., 2016, 2017). To scale to large data sets, minibatches of both cells and genes are used to approximate gradients at each update. Genes (i.e., features) are sampled in proportion to the percentage of cells in which they are expressed to efficiently approximate the covariance matrix calculated during inference, inspired by previous random matrix algorithms to approximate high dimensional matrix multiplication (Drineas et al.). Cells (i.e., samples) are sampled uniformly in every batch. Inference was performed on Microsoft Azure High Performance Computing cores.

### Single cell RNA-seq data

We chose four data sets to evaluate tGPLVM’s applicability to identifying cell type, state, and developmental trajectory and scalability to experiments with large numbers of cells. The Pollen data (Pollen et al.), which were used to evaluate clustering, consist of 11 distinct mouse neural and blood cell populations across 249 cells sequenced on a Fluidigm C1 systems. Pollen is a dense matrix because of high read depth, with about 80% non-zero values. The counts are log normalized as log_10_(1 + *Y*). Inference of development trajectories was evaluated on the data used to develop the method GPfates from Lonnberg (Lönnberg etal., 2017). Lonnberg (Lönnberg etal., 2017) sequenced 408 T helper cells cells over 7 days after *Plasmodium* infection on a Fluidigm C1 system. The Lonnberg data are provided as TPM measurements. The data are sparse and normalized by log_2_(1 + *Y*). Cell state was explored on batch of CD34+ peripheral blood mononuclear cells (Zheng et al., 2017) (PBMCs). About 10,000 cells were captured with 10x Cell Ranger sequencing technologies. These sparse data were also normalized as *log*_2_(1 + *Y*). Finally, to ensure scalability to the most recent experimental data sets, we fit the model to 1 million mice brain cells sequenced on a 10x Cell Ranger (Zheng et al., 2017). We normalized the mice brain cells as log2(1+*Y*)

#### Identifying cell types with k-means clustering

Clustering for cell type identification was evaluated on the Pollen (Pollen et al.) data. tGPLVM and comparison methods were used to fit latent mappings between 2 and 9 dimensions. To perform clustering for each of the estimated latent manifolds, we used k-means clustering with the number of clusters equal to the number of different cell type or cell state labels in the existing data. Clustering with k-means was repeated 10 times on the mean of the posterior of the latent position and evaluated against true labels using normalized mutual information (NMI) and adjusted rand score (ARS). Mutual information measures the amount of information contained in one random variable, the true labels, by another random variable, the inferred labels. NMI normalizes mutual information by the geometric mean of the entropy of both labels to a scale of zero - no mutual information - to one – the same distribution (Strehl and Ghosh, 2003). ARS is a measure of the proportion of shared members between pairs of true and estimated clusters (Hubert and Arabie). Zero inflated factor analysis (ZIFA) (Pierson and Yau), t-SNE (Van Der Maaten and Hinton) (perplexity set to default 30), and PCA (Hotelling, 1933) were tested as comparison methods. To evaluate the robust adaptations of the tGPLVM model, we fit tGPLVM with only an SE kernel or SE and Matérn 1/2 kernel as well as tGPLVM with normally distributed error.

#### Trajectory building with minimum spanning trees

tGPLVM was used to fit a two dimensional latent mappings for the Lonnberg (Lönnberg etal., 2017) developmental data. The minimum spanning tree was found on Euclidean distance matrix of the posterior means of the low dimensional embedding and compared to sequencing time to verify correct ordering. The same analysis was performed with ZIFA (Pierson and Yau), t-SNE (Van Der Maaten and Hinton) (perplexity set to default 30), and PCA (Hotelling, 1933).

#### Visualization of sparse, raw count matrices

tGPLVM was used to fit a three-dimensional mapping for the two 10x data sets, CD34+ cells and mice brain cells. Pearson correlation between latent position posterior mean and expression counts was used to identify genes associated with latent dimensions. ZIFA (Pierson and Yau), t-SNE (Van Der Maaten and Hinton) (perplexity set to default 30), and PCA (truncated SVD (Halko et al.)) were also fit to the CD34+ cell data to compare computational time.

### Scaling inference to high-throughput experiments

Computational times were recorded for samples of 100, 1,000, 10,000, 100,000 and 1,000,000 cells from the 10x 1 million mouse brains cells (Zheng et al., 2017) to a latent embedding using tGPLVM and comparison methods. Experiments were run on a Standard H16m (16 VCPUs, 224 GB memory) Azure high performance computing Unix system. tGPLVM was run for 100 passes through the data. Each minibatch contained the minimum of 2500 or the number of cells and 250 genes. ZIFA, t-SNE, and PCA (using truncated SVD) were fit until convergence or failure.

## Acknowledgments

BEE was funded by CZI AWD1005664, CZI, CZI AWD1005667, NIH R01 HL133218, NIH U01 HG007900, a Sloan Faculty Fellowship, and an NSF CAREER AWD1005627.

The authors declare that they have no competing financial interests.

Correspondence and requests for materials should be addressed to B.E.E.

Software is available at https://github.com/architverma1/tGPLVM

## Author Contributions

A.V. and B.E.E designed experiments, analyzed experiments, and wrote paper. A.V. wrote implementation and ran experiments.

## References

Sumon Ahmed, Magnus Rattray, and Alexis Boukouvalas. Grandprix: Scaling up the bayesian gplvm for single-cell data. Bioinformatics, page bty533, 2018. doi: 10.1093/bioinformatics/bty533.

Matthew Amodio, Krishnan Srinivasan, David van Dijk, Hussein Mohsen, Kristina Yim, Rebecca Muhle, Kevin R Moon, Susan Kaech, Ryan Sowell, Ruth Montgomery, James Noonan, Guy Wolf, and Smita Krishnaswamy. Exploring Single-Cell Data with Multitasking Deep Neural Networks. bioRxiv. doi: 10.1101/237065.

Philipp Angerer, Laleh Haghverdi, Maren Bttner, Fabian J. Theis, Carsten Marr, and Florian Buettner. destiny: diffusion maps for large-scale single-cell data in r. Bioinformatics, 32(8):1241–1243, 2016. doi: 10.1093/bioinformatics/btv715.

J L Browning, A Ngam-ek, P Lawton, J DeMarinis, R Tizard, E P Chow, C Hesslon, B O’Brine-Greco, S Foley, and C F Ware. Lymphotoxin B, a novel member of the TNF family that forms a heteromeric complexs with lymphotoxing on the cell surface. Cell, 72:847–856, 1993.

Florian Buettner, Kedar N Natarajan, F Paolo Casale, Valentina Proserpio, Antonio Scialdone, Fabian J Theis, Sarah A Teichmann, John C Marioni, and Oliver Stegle. Computational analysis of cell-to-cell heterogeneity in single-cell RNA-sequencing data reveals hidden subpopulations of cells. Nature Biotechnology, page 155, jan.

Andrew Butler, Paul Hoffman, Peter Smibert, Efthymia Papalexi, and Rahul Satija. Integrating single-cell transcriptomic data across different conditions, technologies, and species. Nature Biotechnology, page 411, apr.

Pierre Comon. Independent component analysis, A new concept. Signal Processing, 36(3): 287–314, 1994. ISSN 01651684. doi: 10.1016/0165-1684(94)90029-9.

Andreas C Damianou, Michalis K Titsias, and Neil D Lawrence. Variational inference for latent variables and uncertain inputs in Gaussian processes. Journal of Machine Learning Research, 17:1–62, 2016. ISSN 15337928.

R. Donato, B. R. Cannon, G. Sorci, F. Riuzzi, K. Hsu, D. J. Weber, and C. L. Geczy. Functions of S100 Proteins. Current Molecular Medicine, (1):24–57. ISSN 15665240. doi: 10.2174/156652413804486214.

Petros Drineas, Ravi Kannan, and Michael W. Mahoney. Fast Monte Carlo Algorithms for Matrices I: Approximating Matrix Multiplication. SIAM Journal on Computing, (1): 132–157. ISSN 0097-5397. doi: 10.1137/S0097539704442684.

Barbara E. Engelhardt and Matthew Stephens. Analysis of population structure: A unifying framework and novel methods based on sparse factor analysis. PLoS Genetics, 6(9), 2010. ISSN 15537390. doi: 10.1371/journal.pgen.1001117.

Gökcen Eraslan, Lukas M Simon, Maria Mircea, Nikola S Mueller, and Fabian J Theis. Single cell RNA-seq denoising using a deep count autoencoder. bioRxiv, 2018. doi: 10.1101/300681.

Jean Fan, Neeraj Salathia, Rui Liu, Gwendolyn E. Kaeser, Yun C Yung, Joseph L Herman, Fiona Kaper, Jian-Bing Fan, Kun Zhang, Jerols Chun, and Peter V Kharchenko. Characterizing transcriptional heterogeneity through pathway and gene set overdispersion analysis. Nature Methods, 13(13):241–244, 2016. ISSN 1548-7105.

Laleh Haghverdi, Maren Büttner, F. Alexander Wolf, Florian Buettner, and Fabian J. Theis. Diffusion pseudotime robustly reconstructs lineage branching. Nature Methods, 13(10): 845–848, 2016. ISSN 15487105. doi: 10.1038/nmeth.3971.

Nathan Halko, Per-Gunnar Martinsson, and Joel A. Tropp. Finding structure with randomness: Probabilistic algorithms for constructing approximate matrix decompositions. pages 1–74. ISSN 0036-1445. doi: 10.1137/090771806.

Harry H Harman. Modern factor analysis. 1960.

Harold Hotelling. Analysis of a complex of statistical variables into principal components. Journal of educational psychology, 24(6):417, 1933.

Harold Hotelling. Relations between two sets of variates. Biometrika, 28(3-4):321–377, 1936. doi: 10.1093/biomet/28.3-4.321.

Lawrence Hubert and Phipps Arabie. Comparing partitions. Journal of Classification, (1): 193–218, dec. ISSN 1432-1343. doi: 10.1007/BF01908075.

Neil Lawrence. Probabilistic non-linear Principal Component Analysis with Gaussian Process Latent Variable Models. Journal ofMachine Learning Research, pages 1783–1816. ISSN 1476-4687.

Wei Vivian Li and Jingyi Jessica Li. An accurate and robust imputation method scImpute for single-cell RNA-seq data. Nature Communications, 9(1):997, 2018. ISSN 2041-1723. doi: 10.1038/s41467-018-03405-7.

Tapio Lönnberg, Valentine Svensson, Kylie R James, Daniel Fernandez-Ruiz, Ismail Sebina, Ruddy Montandon, Megan S F Soon, Lily G Fogg, Arya Sheela Nair, Urijah N. Liligeto, Michael J T Stubbington, Lam-Ha Ly, Frederik Otzen Bagger, Max Zwiessele, Neil D Lawrence, Fernando Souza-Fonseca-Guimaraes, Patrick T Bunn, Christian R Engwerda, William R Heath, Oliver Billker, Oliver Stegle, Ashraful Haque, and Sarah A Teichmann. Single-cell RNA-seq and computational analysis using temporal mixture modeling resolves T H 1/T FH fate bifurcation in malaria. Science Immunology, 2(9):eaal2192, 2017. ISSN 2470-9468. doi: 10.1126/sciimmunol.aal2192.

Romain Lopez, Jeffrey Regier, Michael B Cole, Michael Jordan, and Nir Yosef. Bayesian Inference for a Generative Model of Transcriptome Profiles from Single-cell RNA Sequencing. bioRxiv. doi: 10.1101/292037.

A. O’Hagan. On outlier rejection phenomena in bayes inference. Journal of the Royal Statistical Society. Series B (Methodological), 41(3):358–367, 1979. ISSN 00359246.

A. O’Hagan. Modelling with heavy tails. I. Bayesian statistics, 3 (Valencia, 1987), Oxford Sci. Publ., pages 345–359. Oxford Univ. Press, New York, 1988.

Emma Pierson and Christopher Yau. ZIFA: Dimensionality reduction for zero-inflated single-cell gene expression analysis. Genome Biology, (1):241. ISSN 1474-760X. doi: 10.1186/s13059-015-0805-z.

Alex A Pollen, Tomasz J Nowakowski, Joe Shuga, Xiaohui Wang, Anne A Leyrat, Jan H Lui, Nianzhen Li, Lukasz Szpankowski, Brian Fowler, Peilin Chen, Naveen Ramalingam, Gang Sun, Myo Thu, Michael Norris, Ronald Lebofsky, Dominique Toppani, Darnell W Kemp II, Michael Wong, Barry Clerkson, Brittnee N Jones, Shiquan Wu, Lawrence Knutsson, Beatriz Alvarado, Jing Wang, Lesley S Weaver, Andrew P May, Robert C Jones, Marc A Unger, Arnold R Kriegstein, and Jay A A West. Low-coverage single-cell mRNA sequencing reveals cellular heterogeneity and activated signaling pathways in developing cerebral cortex. Nature Biotechnology, page 1053, aug.

Rajesh Ranganath, Sean Gerrish, and David M Blei. Black Box Variational Inference. Aistats. ISSN 15337928.

Jaehoon Shin, Daniel A. Berg, Yunhua Zhu, Joseph Y. Shin, Juan Song, Michael A. Bonaguidi, Grigori Enikolopov, David W. Nauen, Kimberly M. Christian, Guo Li Ming, and Hongjun Song. Single-Cell RNA-Seq with Waterfall Reveals Molecular Cascades underlying Adult Neurogenesis. Cell Stem Cell, (3):360–372. ISSN 18759777. doi: 10.1016/j.stem.2015.07.013.

Laura E. Sidney, Matthew J. Branch, Siobhán E. Dunphy, Harminder S. Dua, and Andrew Hopkinson. Concise review: Evidence for CD34 as a common marker for diverse progenitors. Stem Cells, 32(6):1380–1389, 2014. ISSN 15494918. doi: 10.1002/stem.1661.

Gil Stelzer, Naomi Rosen, Inbar Plaschkes, Shahar Zimmerman, Michal Twik, Simon Fishilevich, Tsippi Iny Stein, Ron Nudel, Iris Lieder, Yaron Mazor, Sergey Kaplan, Dvir Dahary, David Warshawsky, Yaron Guan-Golan, Asher Kohn, Noa Rappaport, Marilyn Safran, and Doron Lancet. The GeneCards suite: From gene data mining to disease genome sequence analyses. Current Protocols in Bioinformatics, 2016(June):1.30.1–1.30.33, 2016. ISSN 1934340X. doi: 10.1002/cpbi.5.

Alexander Strehl and Joydeep Ghosh. Cluster ensembles - A knowledge reuse framework for combining multiple partitions. Journal of Machine Learning Research, 3(3):583–617, 2003. ISSN 15324435. doi: 10.1162/153244303321897735.

Qingtao Tang, Li Niu, Yisen Wang, Tao Dai, Wangpeng An, Jianfei Cai, and Shu Tao Xia. Student-t process regression with student-t likelihood. IJCAI International Joint Conference on Artificial Intelligence, pages 2822–2828. ISSN 10450823.

Michalis Titsias and Neil Lawrence. Bayesian Gaussian Process Latent Variable Model. Artificial Intelligence, pages 844–851. ISSN 0899-7667. doi: 10.1162/089976699300016331.

Elena Tomasello and Eric Vivier. KARAP/DAP12/TYROBP: Three names and a multiplicity of biological functions. European Journal of Immunology, 35(6):1670–1677, 2005. ISSN 00142980. doi: 10.1002/eji.200425932.

Dustin Tran, Alp Kucukelbir, Adji B. Dieng, Maja Rudolph, Dawen Liang, and David M. Blei. Edward: A library for probabilistic modeling, inference, and criticism. arXiv preprint arXiv:1610.09787, 2016.

Dustin Tran, Matthew D. Hoffman, Rif A. Saurous, Eugene Brevdo, Kevin Murphy, and David M. Blei. Deep probabilistic programming. In International Conference on Learning Representations, 2017.

Cole Trapnell, Davide Cacchiarelli, and Xiaojie Qiu. Monocle: Cell counting, differential expression, and trajectory analysis for single-cell RNA-Seq experiments. Bioconductor, page 10.

L J P Van Der Maaten and G E Hinton. Visualizing high-dimensional data using t-SNE. Journal of Machine Learning Research, pages 2579–2605. ISSN 1532-4435. doi: 10.1007/s10479-011-0841-3.

Jarno Vanhatalo, Pasi Jylänki, and Aki Vehtari. Gaussian process regression with Student-t likelihood. In Y Bengio, D Schuurmans, J D Lafferty, C K I Williams, and A Culotta, editors. Advances in Neural Information Processing Systems 22, pages 1910–1918. Curran Associates, Inc.

Chang Xia, Zachary Braunstein, Amelia C. Toomey, Jixin Zhong, and Xiaoquan Rao. S100 proteins as an important regulator of macrophage inflammation. Frontiers in Immunology, 8(JAN):1–11, 2018. ISSN 16643224. doi: 10.3389/fimmu.2017.01908.

Grace X Y Zheng, Jessica M Terry, Phillip Belgrader, Paul Ryvkin, Zachary W Bent, Ryan Wilson, Solongo B Ziraldo, Tobias D Wheeler, Geoff P McDermott, Junjie Zhu, Mark T Gregory, Joe Shuga, Luz Montesclaros, Jason G Underwood, Donald A Masquelier, Stefanie Y Nishimura, Michael Schnall-Levin, Paul W Wyatt, Christopher M Hindson, Rajiv Bharadwaj, Alexander Wong, Kevin D Ness, Lan W Beppu, H Joachim Deeg, Christopher McFarland, Keith R Loeb, William J Valente, Nolan G Ericson, Emily A Stevens, Jerald P Radich, Tarjei S Mikkelsen, Benjamin J Hindson, and Jason H Bielas. Massively parallel digital transcriptional profiling of single cells. Nature Communications, 8:14049, jan 2017.

